# A T2T Benchmark Reveals How Reference Choice Shapes Human Genome Interpretation

**DOI:** 10.64898/2026.09.07.749735

**Authors:** Zhuo Huang, Yanan Chu, Yabin Tian, Changjun Shao, Jian Wang, Xueqi Zhang, Jing Chen, Zheng Jia, Lili Li, Jiale Li, Guigao Lin, Kuo Zhang, Stylianos E. Antonarakis, Yu Kang, Jie Huang

## Abstract

The completion of telomere-to-telomere (T2T) human genomes has expanded the accessible landscape of human genetic variation, yet benchmark resources remain limited to conventional high-confidence regions defined by existing reference frameworks. Here, we generated a near-perfect diploid T2T genome (T2T-LIN) from a Chinese individual and established assembly-based truth sets by comparison with T2T-YAO, an ancestry-matched near-perfect T2T reference genome. The benchmark showed a heterozygous/homozygous SNV ratio of ~2, consistent with expectations under Hardy–Weinberg equilibrium, and enabled genome-wide evaluation of reference-dependent biases. We found that reference choice substantially influences genome interpretation: ancestry-matched linear T2T references provided the most faithful representation of individual genomic variation and enabled more accurate genome reconstruction than unmatched linear, diploid and graph-based references. Benchmarking previously inaccessible repetitive and structurally complex regions revealed substantial limitations of current variant callers that were masked by conventional metrics. The T2T-LIN and YAO-LIN benchmarks establish a T2T-era framework for evaluating reference-dependent genome interpretation and variant discovery across nearly the complete human genome.

## Main

The completion of telomere-to-telomere (T2T) human genomes has transformed the landscape of human genetic variation by resolving previously inaccessible repetitive and structurally complex regions^1,2^. The availability of highly polished diploid assemblies from individuals of diverse ancestries^3–5^ now enables comprehensive investigation of variation across nearly complete human genome. Although many repetitive and structurally complex regions remain poorly characterized in population studies, accumulated evidence indicates that variation within these regions, including structural variation, copy-number variation, and challenging medically relevant genes (CMRGs), contributes substantially to human phenotypic diversity ^6–10^. Indeed copy-number variation within ribosomal DNA (rDNA) arrays, among the most challenging regions of the human genome, has been associated with diverse human traits^11^, highlighting the importance of accurately characterizing variation beyond conventionally accessible regions. However, accurate interpretation of this expanded genomic landscape requires not only comprehensive reference genomes but also benchmark frameworks capable of determining whether detected variants faithfully represent individual genomes.

Accurate variant discovery is essential for human genetics, rare disease diagnosis, cancer genomics, population studies, and even human history^12–17^. Variant benchmarks are essential for calibrating variant callers and assessing the reliability of detected variants. Existing benchmarks, particularly the Genome in a Bottle (GIAB) benchmark sets based on the LCL (lymphoblastoid cell line) sample HG002 of Eastern European Ashkenazi Jewish ancestry, have established standardized frameworks for variant-calling evaluation through high-confidence truth sets^18^. Variant truth sets, defined relative to specific reference frameworks, were primarily derived from concordance among sequencing platforms, variant callers and orthogonal evidence. This strategy has been highly effective within accessible high-confidence regions, which encompass ~85% of the human genome, but limits systematic evaluation of variation in unresolved repetitive regions and prevents direct assessment of reference-dependent biases^19–24^. Although T2T genomes have recently expanded the accessible genomic landscape, high-confidence truth sets describing genome-wide variation between complete individual genomes are still needed to enable systematic evaluation of variant discovery across these newly resolved regions^25^. Moreover, existing benchmarks have largely evaluated variant discovery against a single reference representation, leaving unresolved whether reference choice itself systematically influences genome interpretation. With multiple linear T2T references and emerging graph-based representations now available, determining how reference choice and genome representation affect variant discovery and genome reconstruction has become an important challenge.

Here, we generated T2T-LIN, a *de novo* assembled near-perfect diploid T2T genome from a Han Chinese individual, as an East Asian analogue of the widely used HG002 benchmark genome. We used T2T-LIN as an experimentally validated genome benchmark to evaluate variant discovery and personal genome reconstruction across existing reference frameworks, and found that the ancestry-matched T2T-YAO genome^4^ provided the most faithful reconstruction of LIN. We then established assembly-based truth sets through direct comparison of the two complete diploid genomes to construct the YAO-LIN variant benchmark. This framework extends high-confidence benchmarking into previously inaccessible genomic regions and enables systematic evaluation of variant discovery across nearly the complete human genome. Benchmarking current variant callers revealed substantial performance degradation in repetitive and structurally complex regions that are largely excluded from existing benchmarks. Together, these results establish a T2T-era framework for evaluating reference-dependent genome interpretation and variant discovery across complete human genomes.

## Results

### Building T2T-LIN, a near-perfect diploid human genome benchmark

Existing human genome benchmarks have substantial ancestry imbalance, with East Asian populations remaining underrepresented despite their large population size. To extend genome benchmarking beyond conventional accessible regions and enable ancestry-aware evaluation of complete human genomes, we generated a *de novo* assembled near-perfect diploid telomere-to-telomere genome, T2T-LIN, from one member of a male twin pair (QI•LIN) of southern Chinese origin. Analogous to the HG002 v1.1, T2T-LIN provides a genome benchmark for personal genome reconstruction and together with T2T-YAO^4^, enables variant benchmarking across nearly the complete human genome. Importantly, the YAO-LIN benchmark captures observed genome-wide genetic differences between two individuals from the same ethnic group, providing an ancestry-matched framework for evaluating reference-dependent variant discovery. In contrast, the canonical references GRCh38 and CHM13^1^ are not complete diploid assemblies derived from individual genomes and therefore cannot provide an equivalent individual-level framework for genome reconstruction and variant benchmarking.

We chose a lymphoblastoid cell line (LCL) as the benchmark material for LIN as for other widely used benchmark samples^26^. LCLs can accumulate cell-line-specific somatic variation^27^ that should be excluded from benchmarking regions to avoid confounding somatic variation with germline differences when evaluating variant-calling accuracy. However, comprehensive characterization of such variation remains challenging at the scale of a complete diploid T2T genome. For HG002, for example, only 85 somatic SNPs have been reported within the high-confidence benchmark regions^28^, leaving the genome-wide extent of cell-line-associated somatic variation incompletely characterized.

To achieve T2T-LIN near-perfect quality and systematically characterize somatic variation relevant to benchmarking, we generated complementary sequencing data using ultra-long Oxford Nanopore R10, PacBio SPRQ HiFi, Element AVITI, and Pore-C platforms (**Table S1**). Assembly and polishing followed the pipeline developed for T2T-YAO v2.0^4^. The resulting diploid assembly achieved T2T completeness except for immunoglobulin clusters affected by somatic rearrangement^29^. SAS evaluation^4^ provided sufficient support from aligned reads for 99.89% of the diploid T2T-LIN genome, with only 6.63 Mb lacking sufficient alignment support, comprising 4.01 Mb rDNA clusters, 530 long homopolymer tracts, six non-homopolymer sites (<50 bp) outside rDNA clusters, and collapsed sequence totaling 2.62 Mb within chromosome Yq12. The assembly achieved k-mer-based QVs of 76.92 for 21-mers and 68.47 for 31-mers, similar to those reported for other near-perfect human genomes^3,4^.

We further identified 408.33 kb of regions affected by LCL-derived somatic variation^27^, including 21 structural variants totally spanning 402 kb and 5,890 SNVs and indels (<50 bp) identified as somatic events by Sniffles2^30^ and DeepVariant^31,32^, respectively. Together with rDNA arrays and telomeric regions, which may exhibit mitotic instability^33,34^, these regions were excluded from benchmark construction (**Table S2**). Thus, T2T-LIN provides a near-complete, high-quality diploid genome with explicitly characterized regions that could confound genome-wide benchmarking.

### Construction of assembly-based variant truth sets for current references

Variant truth sets are collections of validated variants used for benchmarking variant discovery relative to a specific reference genome. Using T2T-LIN as an experimentally validated genome benchmark, we constructed assembly-based truth sets directly from genome-to-genome comparisons, rather than inferring truth primarily from concordance among sequencing platforms and variant callers. We considered three linear reference frameworks represented by a single haplotype: GRCh38, T2T-CHM13, and T2T-YAO^1,4^. These represent an established pre-T2T reference of mosaic origin (GRCh38), an unmatched T2T reference of European ancestry (CHM13), and an ancestry-matched T2T reference (YAO), respectively. Because T2T-YAO is a diploid assembly, we first evaluated the concordance of its maternal and paternal haplotypes across autosomes. Unalignable regions in the paternal haplotype were mainly restricted to centromeres and acrocentric short arms, totaling 14.94 Mb outside rDNA and telomeric regions, compared with 57.96 Mb in the maternal haplotype, excluding the sex chromosomes. We therefore selected the maternal haplotype, which provides a greater amount of alignable sequence for reads from other samples, together with chromosome Y (YAO-mat+Y, hereafter YAO), as the representative linear reference for subsequent analyses.

T2T-LIN was independently aligned to each reference to identify homozygous and heterozygous variants. Candidate SNVs, indels, and structural variants (SVs) were identified using dipcall^35^, while gene copy-number variants (gCNVs) were determined by identifying additional gene copies (identity >0.95) after lifting annotations from GRCh38 using Liftoff ^36^. To establish high-confidence truth sets, candidate variants were further evaluated using independent sequencing evidence from all three sequencing platforms. We extended the sufficient alignment support framework (SAS)^37^ from assembly validation to variant-level validation (SAS-var), requiring candidate variants to be supported by independent evidence from complementary sequencing technologies (**Fig. 1a; Methods**). Unlike existing truth sets that rely primarily on concordance among sequencing platforms and variant callers^22–24,38^, our benchmark derives candidate variants directly from whole-genome assemblies and subsequently validates each variant using complementary sequencing evidence.

**Fig. 1.**
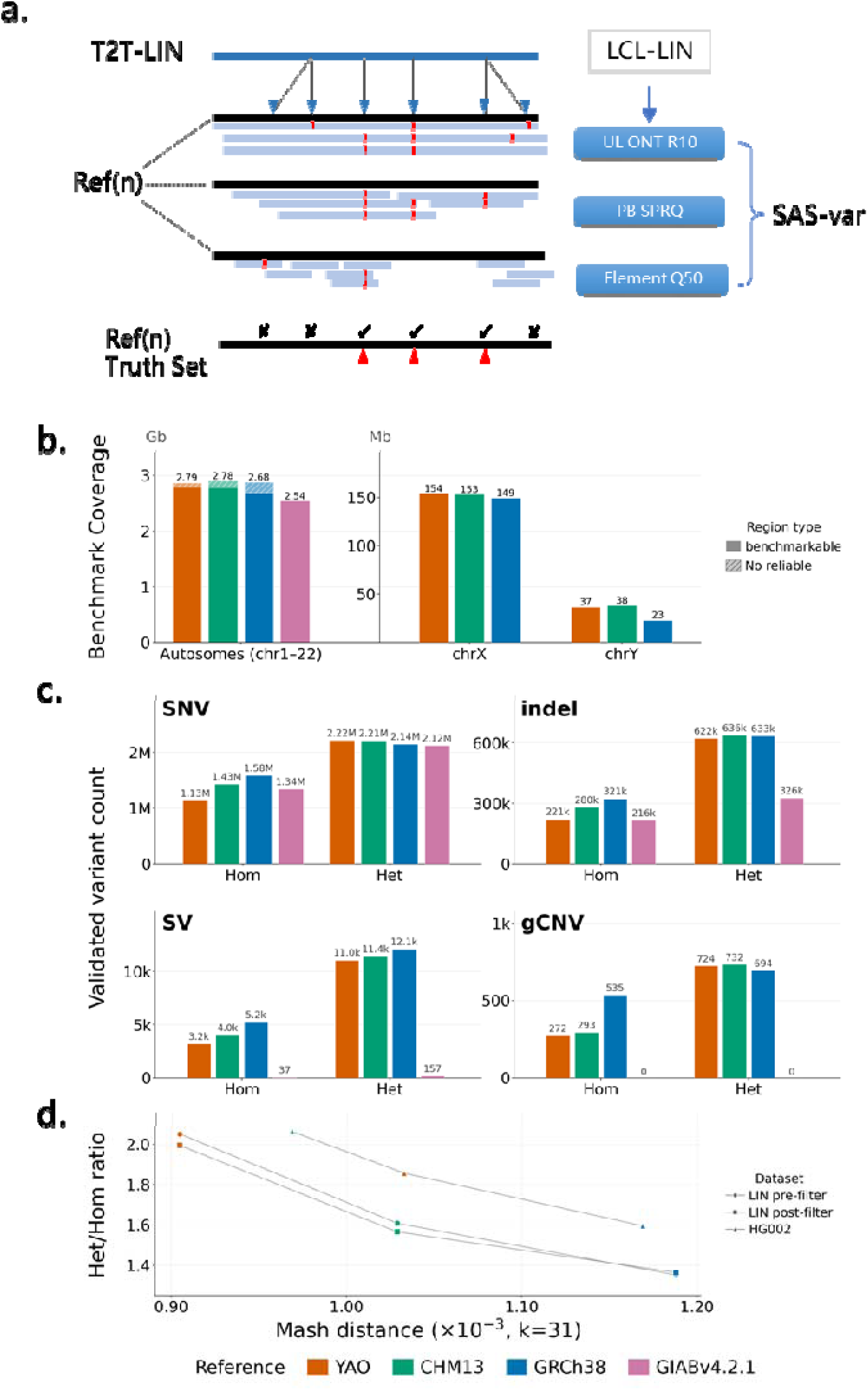
Construction of support-validated, reference-aware benchmark truth sets. **a,** Overview of assembly-based truth-set construction. A support-validated diploid T2T genome (T2T-LIN) was generated from LCL-LIN using Oxford Nanopore ultra-long R10, PacBio SPRQ HiFi, and Element Q50 sequencing. Candidate variants identified by assembly comparison were validated using variant-level Sufficient Alignment Support (SAS-var), generating high-confidence truth sets. **b,** Benchmarkable genomic regions across autosomes, chromosome X, and chromosome Y for YAO–LIN, CHM13–LIN, GRCh38–LIN, and GIAB v4.2.1 benchmarks. Hatched regions indicate excluded intervals lacking reliable truth sets. **c,** Numbers of validated homozygous (Hom) and heterozygous (Het) SNVs, indels, SVs, and gCNVs identified using different reference genomes. **d,** Relationship between reference genetic distance and Het/Hom ratio. Mash distance (31-mers) was used to estimate genetic distance between LIN and each reference. Ratios are shown before and after removal of somatic heterogeneous variants (LIN pre-filter and post-filter), with HG002 shown for comparison. The ancestry-matched YAO reference produced Het/Hom ratios closest to the expected value under Hardy–Weinberg equilibrium.

Relative to GIAB v4.2.1^38^, the assembly-based truth sets substantially expanded *benchmarkable* genomic space (**Table 1**). Across all the various reference sequences, T2T-LIN-based benchmarks increased accessible benchmark regions by approximately 240 Mb in autosomes and 170 Mb in sex chromosomes, with T2T references providing broader coverage than GRCh38 (**Fig. 1b**). The expanded benchmarks contained substantially more validated variants, including SNVs, indels, SV, and gCNV (**Fig. 1c**). Whereas GIAB v4.2.1 contains 194 validated SVs and no benchmarked gCNVs for systematic evaluation, the assembly-based benchmark identified approximately 15,000 SVs and 1,000 gCNVs (**Table S3**), extensively expanded their truth set.

**Table 1.**
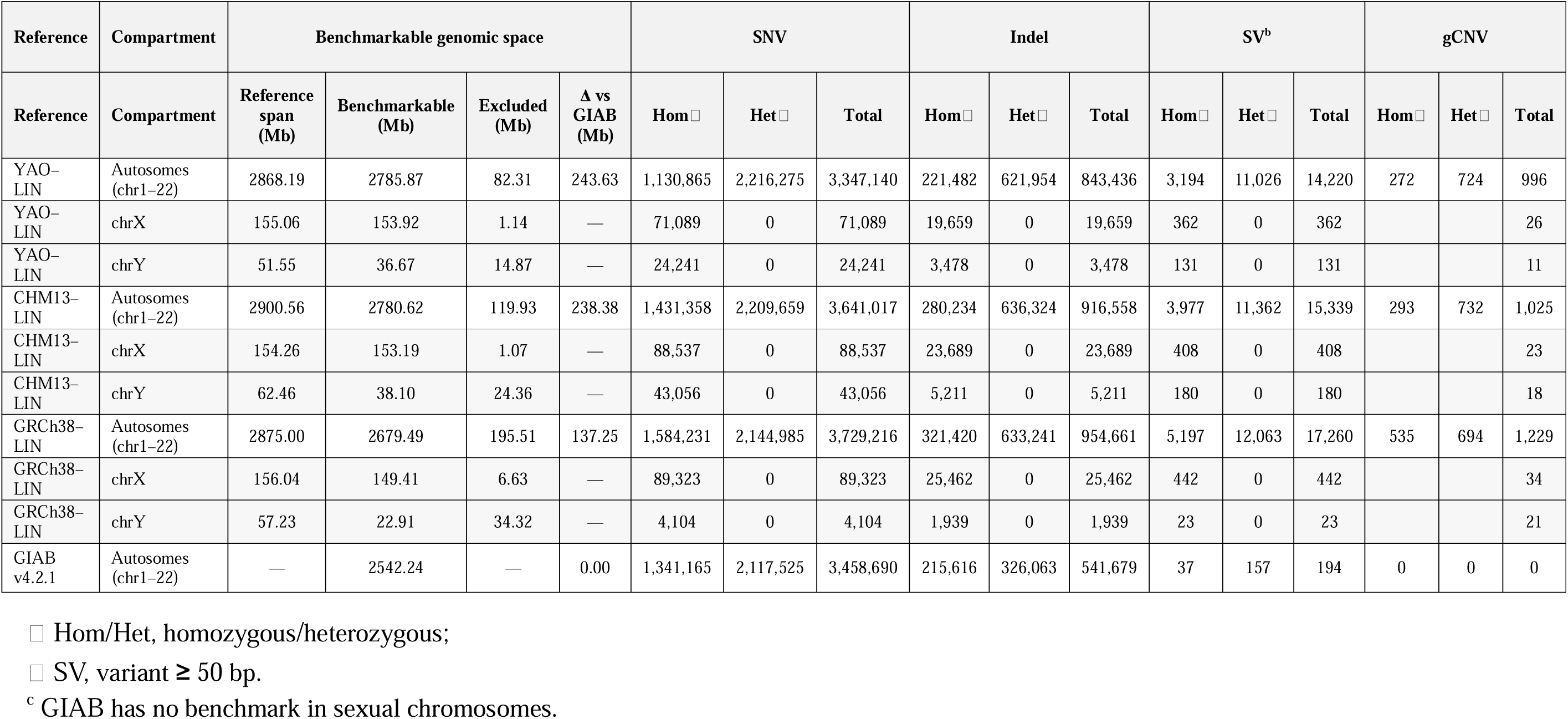
Comparison of T2T-LIN assembly-based truth sets with the GIAB v4.2.1 benchmark.

| Reference | Compartment | Benchmarkable genomic space |  |  |  | SNV |  |  | Indel |  |  | SV<sup>b</sup> |  |  | gCNV |  |  |
| --- | --- | --- | --- | --- | --- | --- | --- | --- | --- | --- | --- | --- | --- | --- | --- | --- | --- |
| Reference | Compartment | Reference span (Mb) | Benchmarkable (Mb) | Excluded (Mb) | Δ vs GIAB (Mb) | Hom ☐ | Het ☐ | Total | Hom ☐ | Het ☐ | Total | Hom ☐ | Het ☐ | Total | Hom ☐ | Het ☐ | Total |
| YAO-LIN | Autosomes (chr1–22) | 2868.19 | 2785.87 | 82.31 | 243.63 | 1,130,865 | 2,216,275 | 3,347,140 | 221,482 | 621,954 | 843,436 | 3,194 | 11,026 | 14,220 | 272 | 724 | 996 |
| YAO-LIN | chrX | 155.06 | 153.92 | 1.14 | — | 71,089 | 0 | 71,089 | 19,659 | 0 | 19,659 | 362 | 0 | 362 |  |  | 26 |
| YAO-LIN | chrY | 51.55 | 36.67 | 14.87 | — | 24,241 | 0 | 24,241 | 3,478 | 0 | 3,478 | 131 | 0 | 131 |  |  | 11 |
| CHM13-LIN | Autosomes (chr1–22) | 2900.56 | 2780.62 | 119.93 | 238.38 | 1,431,358 | 2,209,659 | 3,641,017 | 280,234 | 636,324 | 916,558 | 3,977 | 11,362 | 15,339 | 293 | 732 | 1,025 |
| CHM13-LIN | chrX | 154.26 | 153.19 | 1.07 | — | 88,537 | 0 | 88,537 | 23,689 | 0 | 23,689 | 408 | 0 | 408 |  |  | 23 |
| CHM13-LIN | chrY | 62.46 | 38.10 | 24.36 | — | 43,056 | 0 | 43,056 | 5,211 | 0 | 5,211 | 180 | 0 | 180 |  |  | 18 |
| GRCh38-LIN | Autosomes (chr1–22) | 2875.00 | 2679.49 | 195.51 | 137.25 | 1,584,231 | 2,144,985 | 3,729,216 | 321,420 | 633,241 | 954,661 | 5,197 | 12,063 | 17,260 | 535 | 694 | 1,229 |
| GRCh38-LIN | chrX | 156.04 | 149.41 | 6.63 | — | 89,323 | 0 | 89,323 | 25,462 | 0 | 25,462 | 442 | 0 | 442 |  |  | 34 |
| GRCh38-LIN | chrY | 57.23 | 22.91 | 34.32 | — | 4,104 | 0 | 4,104 | 1,939 | 0 | 1,939 | 23 | 0 | 23 |  |  | 21 |
| GIAB v4.2.1 | Autosomes (chr1–22) | — | 2542.24 | — | 0.00 | 1,341,165 | 2,117,525 | 3,458,690 | 215,616 | 326,063 | 541,679 | 37 | 157 | 194 | 0 | 0 | 0 |

☐ Hom/Het, homozygous/heterozygous; ☐ SV, variant ≥ 50 bp. <sup>c</sup> GIAB has no benchmark in sexual chromosomes.

We next examined how reference representation influences the composition of benchmark variants across autosomes. Reference selection substantially influenced the variant identification: as the reference became genetically more distant from LIN, homozygous variants increased whereas the heterozygous-to-homozygous (Het/Hom) ratio progressively decreased. Under Hardy–Weinberg equilibrium, the expected Het/Hom ratio is approximately 2 (see Supplementary Note for demonstration). After excluding potential somatic variants (supporting read ratio <0.2), the Het/Hom ratio approached the expected value for YAO (1.99) but decreased substantially for CHM13 (1.57) and GRCh38 (1.36). Similar patterns were observed for HG002 benchmarked against matched CHM13 (2.06), unmatched YAO (1.86), and GRCh38 (1.58), with the Het/Hom ratio showing a negative correlation with the genetic distance between the benchmark genome and the reference genome (**Fig. 1d**). The ratio observed with GRCh38 agrees with previous population studies ^39–41^ while the overall trend indicates that ancestry-matched references minimize reference bias by preserving the expected representation of heterozygous and homozygous variation. Together, these results demonstrate that reference representation is not merely a coordinate choice but fundamentally shapes the observed landscape of human genetic variation. The support-validated T2T-LIN genome therefore provides a reliable framework for quantifying reference bias and benchmarking variant discovery across complete human genomes.

### Reference representation shapes variant discovery and genome reconstruction

To investigate how reference representation influences variant discovery and the resulting reconstruction of an individual’s genome, we inferred virtual T2T-LIN genomes from multiple reference frameworks and compared them with the support-validated diploid T2T-LIN assembly. In addition to the three linear references (GRCh38, T2T-CHM13, and T2T-YAO), representing genome sequences organized in a conventional single linear coordinate system, we evaluated four additional representations: (1) the diploid T2T-LIN assembly as a zero-variant control, (2) the haploid LIN reference (LIN-mat+Y) representing a genome differing only by heterozygous variants, (3) the diploid T2T-YAO assembly (YAO-dip), and (4) the HPRC v1.1 pangenome^42^. The first two references were used to assess the accuracy of variant discovery against self-references in haploid and diploid forms, whereas the latter two enabled comparison between linear and graph-based reference frameworks. To place these references into a common evolutionary context, we constructed an autosomal k-mer-based phylogeny using all reference haplotypes together with the near-perfect HG002 v1.1 assembly, one of the high-quality assemblies included in HPRC v1.1 (**Fig. 2a**). As expected, YAO was genetically closest to LIN among the linear references, whereas CHM13 and GRCh38 were progressively more distant. The relatively large distance between LIN and GRCh38 likely reflects both greater evolutionary divergence and the incompleteness of the pre-T2T reference assembly.

**Fig. 2.**
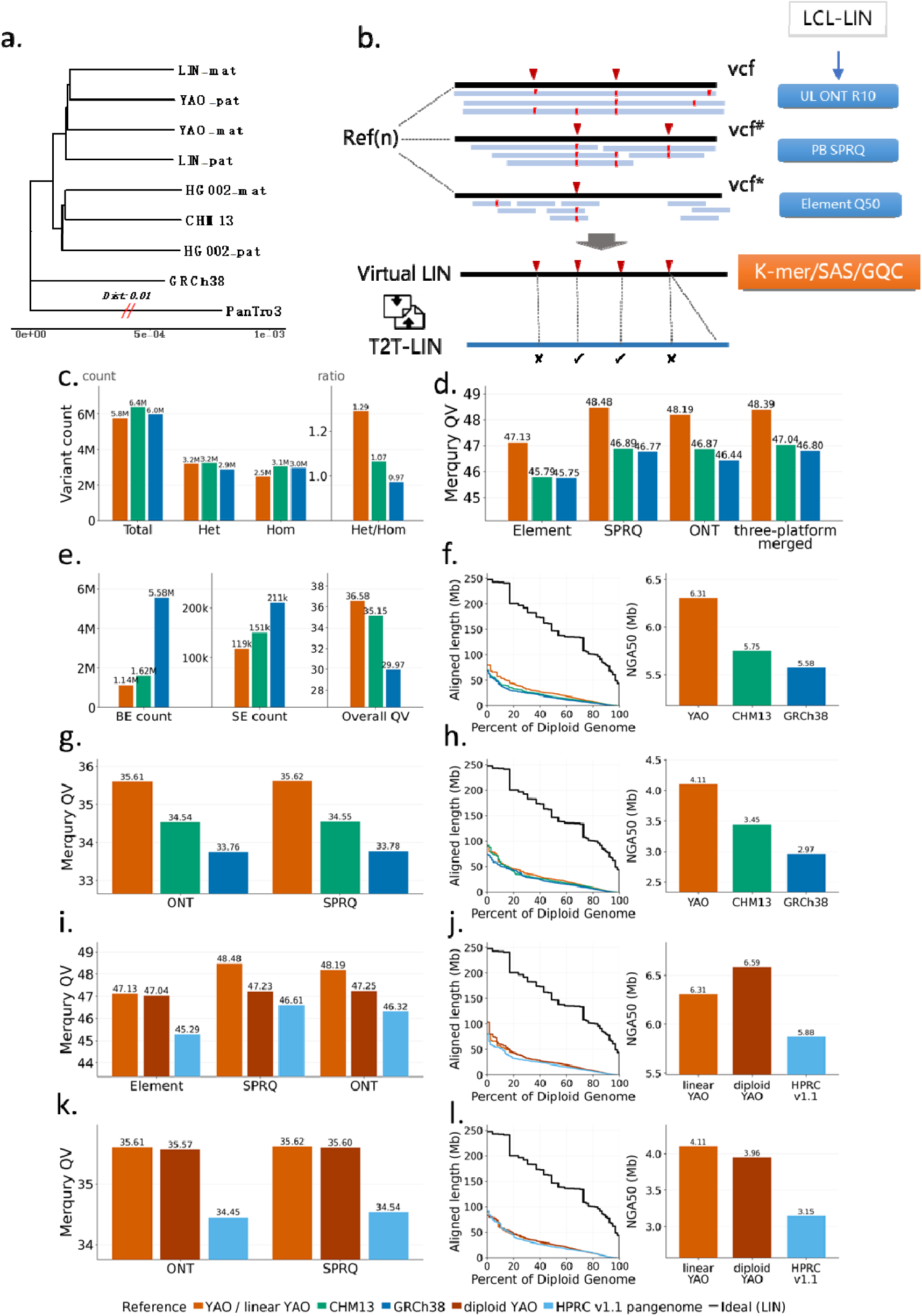
Reference representation shapes variant discovery and reconstruction of personal genomes. **a,** Autosomal k-mer phylogeny of reference haplotypes and representative HPRC genomes. Genetic distance separates ancestry-matched YAO/LIN genomes from CHM13 and GRCh38. **b,** Framework for evaluating genome reconstruction. Variants identified relative to each reference were incorporated to generate virtual diploid LIN genomes, which were assessed against the authentic T2T-LIN assembly using k-mer-, read-, and alignment-based metrics (Merqury, SAS, and GQC). **c,** Numbers and Het/Hom ratios of small variants identified against GRCh38, CHM13, and YAO references. Reference proximity reduces reference-dependent distortion of genotype representation. **d–f,** Evaluation of virtual genomes reconstructed from small variants using Merqury QV (d), SAS-unsupported regions (BE, base-level error; SE, structural error) and SAS-QV(e), and GQC alignment metrics (f). **g–h,** Reconstruction of virtual genomes using structural variants identified against GRCh38, CHM13, and YAO, evaluated by Merqury QV (g) and GQC alignment metrics (h). **i–l,** Evaluation of graph-based references. Small-variant reconstruction using diploid YAO and HPRC v1.1 pangenome representations (i,j), and structural-variant reconstruction (k,l), compared with linear YAO, CHM13, and GRCh38 references. Colors indicate reference representations: YAO/linear YAO (orange), CHM13 (green), GRCh38 (blue), diploid YAO (brown), HPRC v1.1 pangenome (light blue), and ideal T2T-LIN assembly (black).

Using the validated diploid T2T-LIN assembly as the reference, no variants were detected within the benchmarkable regions (**Table S2**), confirming the high accuracy of the assembly. Likewise, when the haploid LIN reference was used, nearly all confidently identified variants were heterozygous and their numbers closely matched direct haplotype-to-haplotype comparisons. Genetic variants are therefore defined and represented as differences between an individual genome and a chosen reference framework, making their discovery inherently reference-dependent. Consequently, an individual’s genome can be either represented as a collection of variants relative to a reference or reconstructed by incorporating those variants into the reference sequence. The correctness of the reconstructed virtual diploid LIN genome can be evaluated by complementary k-mer-, read-, and assembly-based metrics, where the assembly-based metrics using the *de novo* assembled T2T-LIN as the ground truth (**Fig. 2b**). In this framework, T2T-LIN served as a genome benchmark to quantify the fidelity of the entire genome reconstruction process, including sequencing, reference selection, alignment, and variant calling. To facilitate interpretation of homozygous and heterozygous variation, subsequent analyses were restricted to autosomes.

Small variants (SNPs and indels <50 bp) were identified against GRCh38, CHM13, and YAO using DeepVariant^43^ with default parameters. As expected, variant discovery across these three references exhibited a strong dependence on the k-mer-based genetic distance from LIN and reference completeness (**Fig. 2a**). The YAO reference, being genetically closest to LIN, produced the fewest homozygous variants, whereas the Het/Hom ratio progressively declined from YAO to CHM13 and GRCh38 (**Fig. 2c**). We next reconstructed virtual diploid LIN genomes by incorporating the detected variants into each reference and assessed their quality using Merqury (k-mer-based)^44^, SAS (read-based)^37^, and GQC (genome alignment-based)^45^ (**Methods**). Virtual genomes reconstructed from YAO achieved the highest Merqury quality value (QV = 48.48), compared with QV = 47.04 for CHM13 and QV = 46.80 for GRCh38 (**Fig. 2d**), where the sequencing data were used to construct the k-mer sets. Similarly, SAS identified the fewest unsupported windows in the YAO-derived genome, whereas unsupported regions increased by 39% and 4.6-fold in the CHM13- and GRCh38-derived genomes, respectively (**Fig. 2e**). Finally, GQC metric, using the T2T-LIN as ground truth, showed that the YAO-derived virtual genome exhibited the greatest alignment continuity (**Fig. 2f**) and the fewest base-level errors, particularly within homopolymer tracts (**Table S4**). Together, these results indicate that the accuracy of genome reconstruction is strongly influenced by the genetic similarity between the reference and the target genome. Notably, despite extensive optimization of DeepVariant for GRCh38 and its associated benchmark datasets, the larger genetic distance between GRCh38 and LIN still resulted in the greatest reconstruction error. Because Merqury and GQC assessments showed consistent trends with SAS, they were used for evaluating subsequent virtual genomes without requiring read realignment.

To determine whether this finding of least bias in matched reference also applies to structural variation, we identified structural variants relative to each linear reference using Sniffles2^30^ and reconstructed corresponding virtual LIN genomes by incorporating the detected structural variants. The resulting Merqury quality values showed the same trend observed for small variants, although absolute QV values were lower because only structural variants, rather than both small and structural variants, were incorporated (**Fig. 2g**). Consistently, the YAO-derived virtual genome achieved the highest k-mer completeness and the greatest alignment coverage against the authentic T2T-LIN assembly (97.19%), compared with 96.25% for CHM13 and 94.28% for GRCh38 in GQC analysis (**Fig. 2h**). These findings demonstrate that reference choice substantially influences both small-variant and structural-variant discovery and consequently determines the fidelity with which an individual’s genome can be reconstructed.

Finally, we evaluated whether graphic reference with increased diversity could further improve genome reconstruction by comparing the ancestry-matched linear YAO reference with two graph-based references, the diploid YAO assembly (YAO-dip) and the HPRC v1.1 pangenome. Direct reconstruction of a complete individual genome from graph paths generated only ~2 Gb haplotypes using both HPRC v1.1^42^ and CPC^46^, indicating that a substantially larger collection of high-quality haplotypes is required to reconstruct a genome of standard diploid length ^47^. We therefore identified variants through graph-based alignment and projected them onto the primary backbone haplotype (CHM13 for HPRC and YAO-mat for YAO-dip) using the surjection model implemented in the vg pangenome toolkit^48–50^. These variants were then incorporated into the corresponding backbone haplotypes to reconstruct virtual LIN genomes for evaluation. For small variants, none of the graph-based representation improved genome reconstruction relative to the linear YAO reference. Virtual genomes reconstructed from YAO-dip and the HPRC pangenome showed lower Merqury QV values and reduced GQC alignment quality (**Fig. 2i,j**). We next evaluated structural variants using the same framework. Incorporating graph-derived SVs largely restored reconstruction quality for YAO-dip, producing k-mer- and GQC-based metrics comparable to those obtained with the linear YAO reference, whereas the HPRC-derived virtual genome achieved quality similar to that reconstructed from linear CHM13 and consistently exceeded that of GRCh38 (**Fig. 2k,l**). These observations indicate that increasing haplotype diversity alone does not automatically improve reconstruction of complete individual genomes. Performance depends on multiple factors, including backbone reference quality, haplotype completeness, graph alignment accuracy, and fidelity of transferring haplotype variants to linear backbone coordinates. Thus, while pangenomes provide an essential framework for representing human diversity, achieving accurate reconstruction of complete personal genomes will require both comprehensive haplotype resources and improved methods that can accurately infer variants based on graph-reference.

### Current variant-calling performance across T2T genomic regions

The preceding analyses identified the ancestry-matched linear YAO reference as the most suitable framework for benchmark construction in this context, as it provided the most faithful reconstruction of the LIN genome while maintaining substantially lower computational complexity than graph-based approaches. These results highlight that ancestry-matched references improve accuracy when available, and reference selection should account for genetic distance between reference and target genomes. We therefore established the YAO–LIN benchmark and evaluated state-of-the-art SNV, indel, SV, and CNV callers using whole-genome sequencing data generated from LIN by Oxford Nanopore R10, PacBio SPRQ HiFi, and Element platforms. Variant-calling performance was assessed against the YAO reference across genomic stratifications following GA4GH/GIAB benchmarking standards ^23,51^ (**Fig. 3a; Tables S5–S7**). Unlike previous benchmarks, YAO–LIN extends evaluation into complete T2T genomic regions including centromeres, satellite arrays, acrocentric short arms, complete segmental duplications, and multicopy gene families.

**Fig. 3.**
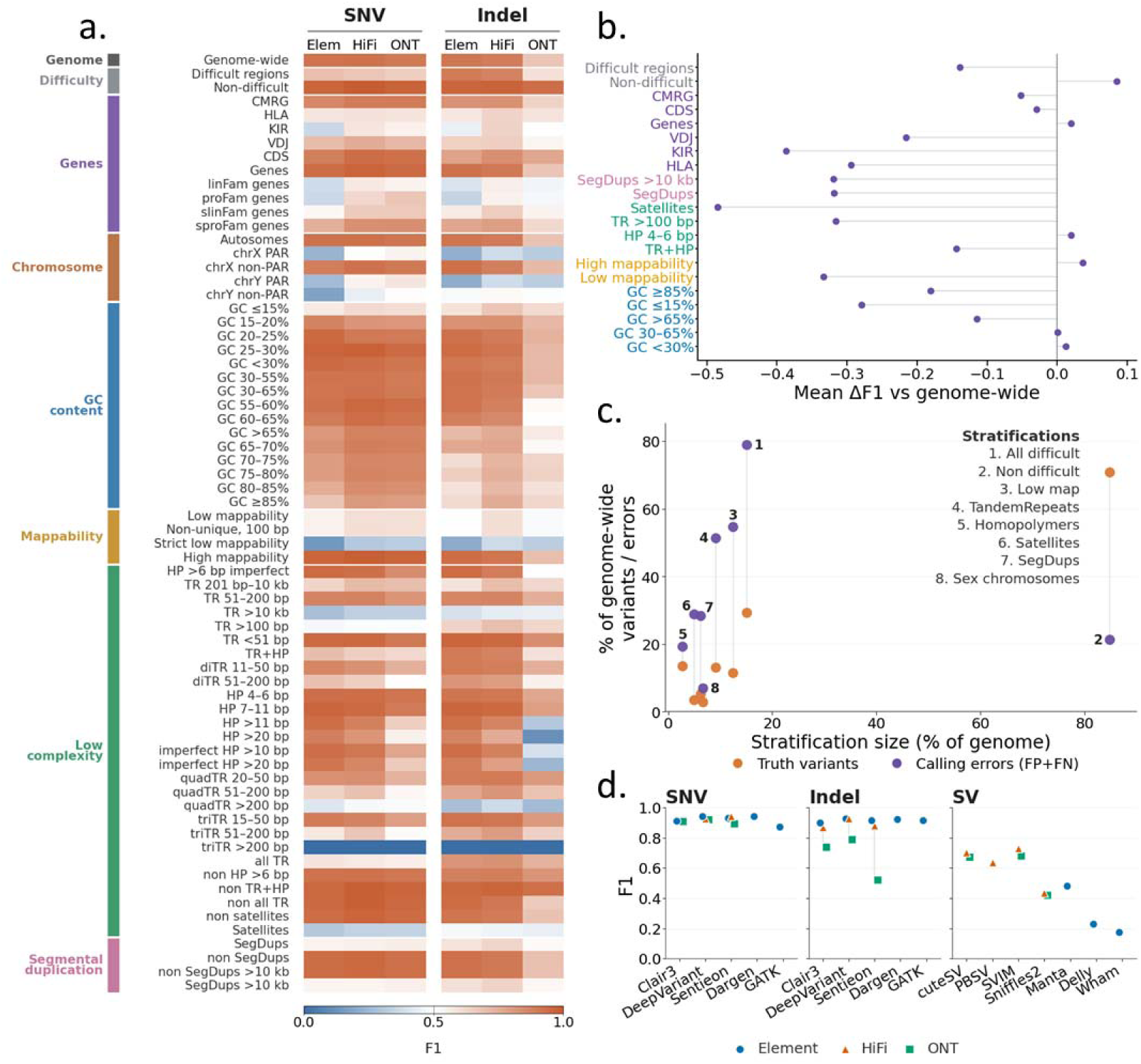
Benchmarking variant-calling performance across complete human genomes using the YAO–LIN benchmark. **a,** Heatmap showing F1 scores of SNV and indel calling workflows across genomic stratifications defined by GA4GH/GIAB standards, including genome-wide, difficult regions, genes, chromosomes, GC content, mappability, low-complexity regions, and segmental duplications. Element (Elem), PacBio HiFi (HiFi), and Oxford Nanopore (ONT) datasets were evaluated independently. **b,** Reduction in F1 score (ΔF1) relative to genome-wide performance across genomic contexts. Complex regions, including satellite DNA, segmental duplications, tandem repeats, and low-mappability regions, show the greatest performance declines. **c,** Contribution of each genomic stratification to truth variants and calling errors. Difficult regions account for a disproportionate fraction of false calls relative to their genomic span. **d,** Performance comparison of representative variant-calling workflows for SNVs, indels, and structural variants across sequencing platforms.

At the whole-genome level, the best-performing workflows achieved F1 scores approaching 0.94 for SNVs and 0.93 for small indels, a little lower than previous report in high-confidence (**Fig. 3a**). However, these relatively excellent genome-wide metrics masked pronounced performance heterogeneity across genomic regions, since the F1 scores are not homogeneous (similar) among these different genomic regions. While many not-difficult regions were effectively saturated, performance deteriorated sharply within repetitive and structurally complex loci, including segmental duplications and multicopy genes, long tandem repeats and satellite-rich regions, low mappability regions and other T2T-enabled genomic compartments. In several of these regions, even the best-performing workflows achieved F1 reductions of −0.1~−0.7 relative to their genome-wide averages (**Fig. 3b**). Although these regions represent a relatively small fraction of the benchmark space, errors within them contributed disproportionately to the total false-positive and false-negative calls (**Fig. 3c**). Consequently, improving variant-calling algorithms and analytical methods for these regions are likely to yield substantially greater gains in variant discovery than further optimization of already well-resolved genomic compartments. Performance also depended strongly on sequencing technology and variant class. Element and HiFi achieved similarly high accuracy for small-variant detection, whereas Oxford Nanopore R10 and HiFi showed comparable performance for structural variant discovery. (**Fig. 3d**). Together, these analyses demonstrate that excellent genome-wide performance can mask substantial underperformance in variant calling within difficult but biologically important regions. The YAO–LIN benchmark therefore provides a more stringent and comprehensive baseline of current variant-calling performance across complete human genomes.

### Complete genome benchmarking reveals hidden limitations of variant discovery

To determine how the YAO–LIN benchmark extends existing benchmarking resources, we compared its genomic coverage with GIAB v4.2.1. Nearly all 2.54 Gb of GIAB benchmark regions could be projected onto the YAO reference. Only 4.39 Mb (0.17%) could not be directly transferred, including 3.04 Mb of unalignable regions and 1.35 Mb of multi-aligned regions, which were enriched for segmental duplications (~43%), low-mappability loci (~41%), MHC regions, and other highly polymorphic sequences. The YAO–LIN benchmark further recovered nearly all projected GIAB regions, excluding only 1.7 Mb (0.07%) of highly polymorphic sequences. Overall, the YAO–LIN benchmark expanded autosomal benchmarkable regions by approximately 10% compared with GIAB. Importantly, these newly accessible regions were highly enriched for genomic regions historically inaccessible to systematic evaluation, including segmental duplications, multicopy genes, satellite arrays, acrocentric short arms, centromeres, and other sequences resolved only after completion of T2T genome assemblies (**Fig.4, Table S8**). These regions contain large numbers of validated variants previously unavailable for systematic assessment. Genome-wide visualization further demonstrated that newly benchmarked regions frequently coincided with sharp declines in variant-calling accuracy (**Fig. 4**), suggesting that previous benchmarks largely excluded the genomic intervals where current methods perform sub-optimally.

**Figure 4.**
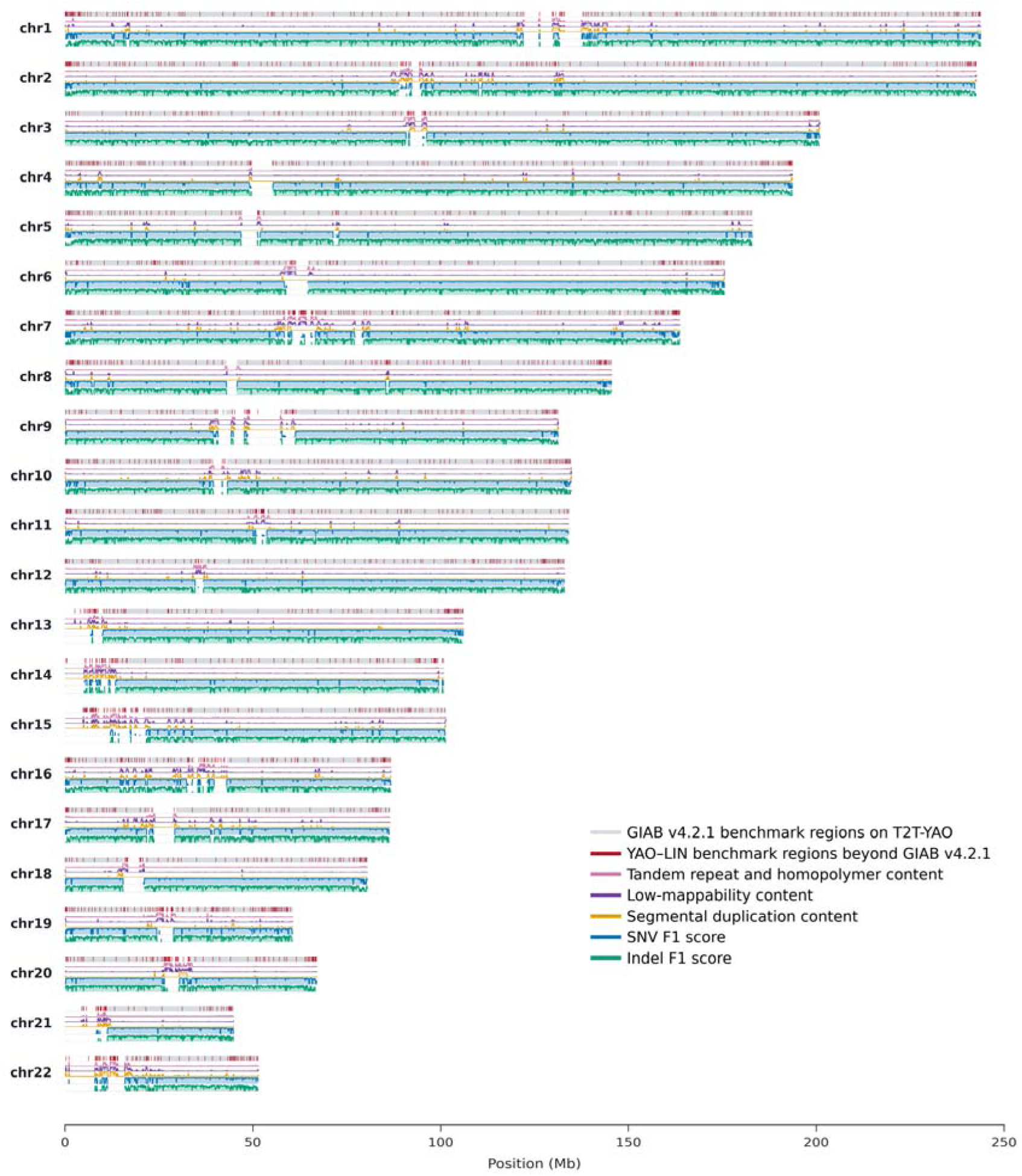
Genome-wide expansion of the YAO–LIN benchmark. Genome-wide distribution of benchmark regions, genomic complexity, and variant-calling accuracy across the autosomes of the YAO reference. Tracks show projected GIAB v4.2.1 benchmark regions (gray), YAO–LIN benchmark extension (red), tandem-repeat and homopolymer content (pink), low-mappability content (purple), segmental duplication content (orange), and F1 scores for SNVs (blue) and indels (green), calculated in 100-kb sliding windows.

To determine whether benchmark expansion reveals previously hidden weaknesses of current variant callers, we divided the benchmark into two categories: regions already represented in GIAB and regions uniquely contributed by the YAO–LIN benchmark (~15% genome size). We found that YAO–LIN extensions were strongly enriched for difficult genomic contexts in both benchmarkable sequence length and number of validated variants, particularly for SVs and gCNVs (**Fig. 5a,b**). Thus, the regions newly established by T2T genomes represent genomic compartments containing substantial amounts of previously inaccessible variation.

**Figure 5.**
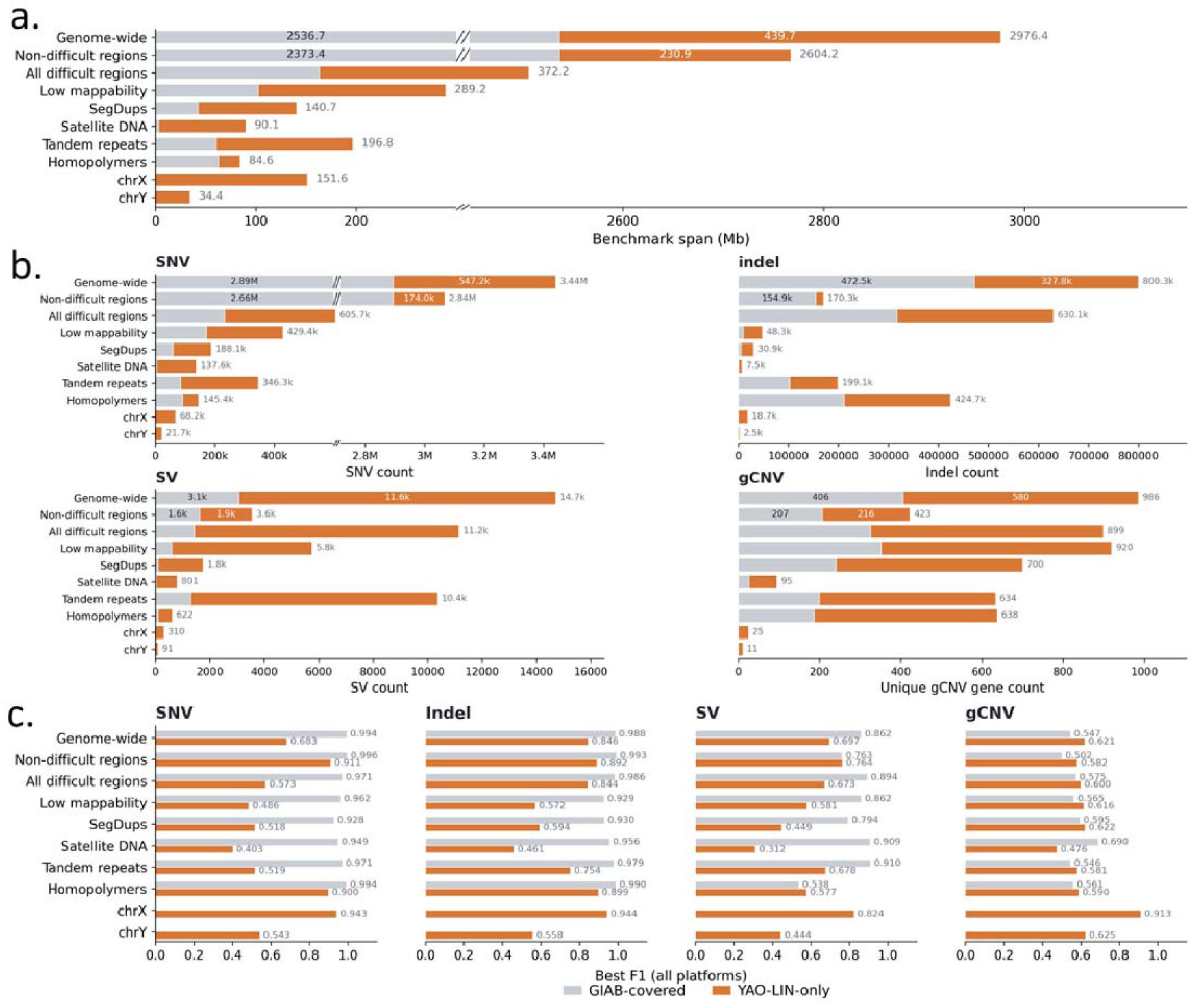
Benchmark expansion reveals previously hidden limitations of variant discovery. **a,** Comparison of benchmark coverage between regions represented by the projected GIAB v4.2.1 benchmark (gray) and regions uniquely contributed by the YAO–LIN benchmark (orange) across representative genomic stratifications. **b,** Numbers of validated single-nucleotide variants (SNVs), insertions/deletions (indels), structural variants (SVs), and genes affected by copy-number variation (gCNVs) contained within GIAB-covered regions and the YAO–LIN benchmark extension. **c,** Best F1 scores achieved by state-of-the-art SNV, indel, SV, and gCNV calling workflows in GIAB-covered regions and the YAO–LIN benchmark extension across representative genomic stratifications.

Comparing the performance of the best-performing workflows for small variants genome-wide and within each stratification, we observed that variant-calling accuracy remained consistently high in non-difficult regions, particularly those represented in GIAB, but declined substantially in difficult regions, especially those newly enabled by the YAO–LIN benchmark (**Fig. 5c**). Consequently, existing benchmarks systematically overestimate genome-wide variant-calling accuracy because they largely exclude the genomic regions where current methods of the pre-T2T era, fail most severely. For SVs and gCNVs, state-of-the-art callers showed limited performance even within GIAB and YAO-LIN regions, highlighting persistent challenges in these variant classes. Collectively, these analyses demonstrate that complete T2T genomes have fundamentally shifted the frontier of human variant benchmarking. By extending systematic evaluation into these regions, the YAO–LIN benchmark reveals the true landscape of current variant-calling performance and provides a roadmap for the development of next-generation variant-calling algorithms.

## Discussion

The availability of complete telomere-to-telomere (T2T) human genomes has expanded the scope of variant discovery, requiring benchmarking frameworks to advance beyond historically accessible genomic regions. Here, we established the YAO-LIN benchmark by directly comparing two independently assembled diploid T2T genomes from ancestry-matched individuals. This assembly-based strategy provides experimentally validated truth sets spanning nearly the complete human genome, overcoming the dependence of previous benchmarks on concordance among sequencing platforms and variant callers. This work demonstrates that major limitations in human genome interpretation are concentrated not in conventional genomic regions, but in repetitive and structurally complex regions that were historically excluded from benchmarking. By extending systematic evaluation into these regions, the YAO–LIN benchmark provides a more realistic assessment of current variant-calling performance and reveals limitations hidden by genome-wide summary metrics. Beyond variant benchmarking, the experimentally validated diploid genomes, accompanied by corresponding benchmark material, the cell line, multi-platform sequencing datasets, YAO-LIN framework, and assembly-based variant truth sets, provide a foundation for evaluating emerging technologies and computational approaches for complete genome analysis^37,52–56^, particularly in highly repetitive and variable regions such as MHC^57–59^, centromeres^60–62^, and other structurally complex genomic compartments. Such resources will become increasingly important as genomic technologies rapidly advance toward scalable and accurate analysis of haplotype-resolved diploid human genomes^47,63–66^.

A major finding of this study is that reference choice substantially influences variant discovery and genome reconstruction. Under current analytical frameworks, ancestry-matched linear T2T references minimized reference bias, preserved expected genotype distributions, and enabled the most faithful reconstruction of individual genomes. These observations highlight a fundamental principle of genome analysis: variants are representations of differences between an individual genome and a selected reference framework. Although graph-based pangenome representations provide a powerful framework for capturing human diversity^67–70^, our results indicate that increasing haplotype diversity alone does not guarantee improved genome reconstruction under current analytical approaches. Realizing the full potential of pangenome-based analysis will require further advances in graph construction, haplotype completeness, graph-native alignment, and variant representation. At present, experimentally validated ancestry-matched T2T references provide a practical complementary strategy for improving accurate variant discovery in complete human genomes^71^. These findings suggest that future genome interpretation frameworks should incorporate both population-aware reference selection and improved graph-based representations to minimize systematic bias.

As complete T2T genomes from diverse populations continue to emerge, experimentally validated and reference-aware benchmarks will be essential for developing next-generation algorithms and population-aware frameworks that accurately characterize human genetic variation across the full complexity of the genome. A future generation of population- and region-specific benchmark genomes and truth sets, including those representing African, South American, and other historically underrepresented populations, will be critical for ensuring that genome analysis methods are evaluated fairly across human genetic diversity.

## Data and code availability

The raw sequencing data for T2T-LIN generated in this study have been deposited in the GSA for Human database of the China National Center for Bioinformation (CNCB) under accession number HRA013169 (Biosample names: CNGB030641(mother), CNGB030644(father), CNGB030643(son, LIN), and are publicly accessible at https://ngdc.cncb.ac.cn/gsa-human. The T2T-LIN genome assemblies are available in the Genome Warehouse at CNCB (https://ngdc.cncb.ac.cn/gwh) under the following accession numbers: GWHJJFJ00000000.1 (maternal), GWHJJFK00000000.1 (paternal).

Code for all workflows are available at https://github.com/KANGYUlab/YAO-LIN-benchmark-workflows.

## Acknowledgments

Lymphoblastoid cell line (LCL-LIN) used for the T2T-LIN assembly were obtained from China National Institute for Drug Control, National Medical Products Administration (https://english.nmpa.gov.cn/2019-07/19/c_389166.htm). The collection of original cells used to establish LCLs was approved by the Institutional Review Board of BGI, China (Approval No. BGI-IRB 22158-T1). The collection and storage of human samples were registered with and approved by the Human Genetic Resources Administration of China (HGRAC). Written informed consents were obtained from the participants. During the preparation of this work, the authors used Deepseek and chatGPT in order to improve language fluency, clarity, and readability. After using this tool or service, the authors reviewed and edited the content as needed and assume full responsibility for the content of the publication.

## Funding

This study was supported by the grants of the National Key Research and Development Program of China (2024YFC3405701 and 2022YFC3400304), the National Science Foundation of China (32371537), National High Level Hospital Clinical Research Funding (BJ-2025-177), S.E.A. is partially supported by the ChildCare Foundation.

## Author contributions

Y.K. and J.H. conceived and designed the project. Z.H. established the YAO-LIN benchmark.

Y.C. conducted the assembly of T2T-LIN. Y.T., C.S., J.W., L.L., and J.L. performed sample preparation and sequencing. Z.H., X.Z., J.C., Z.J., and G.L. participated in data analysis and data presentation. Y.K., J.H., S.E.A., and K.Z. wrote the manuscript. All authors have read and approved the manuscript.

## Declaration of interests

All authors declare no competing interests.

## Supplementary Information

Supplementary text

Table S1 | Sequencing data used for T2T-LIN diploid assembly

Table S2 | Regions and variants excluded from the T2T-LIN truth set

Table S3 | YAO-LIN gene copy-number variants

Table S4 | GQC summary for all virtual / mapped diploid queries

Table S5 | Genomic stratifications used for YAO–LIN benchmark evaluation

Table S6 | Software, versions and models used for YAO–LIN variant benchmarking

Table S7 | Stratified SNV performance by sequencing platform (hap.py)

Table S8 | Expansion of the YAO-LIN benchmark relative to GIAB v4.2.1 by genomic stratification

## Notes

### Competing Interest Statement

The authors have declared no competing interest.

