## Supplemental Note for "A T2T Benchmark Reveals How Reference Choice Shapes Human Genome Interpretation"

SUPPLEMENTARY NOTE

### Derivation of the Heterozygous-to-Homozygous Ratio in Diploid Genome Comparisons

*Statements and proofs*


Let $Q_1, Q_2$ be the two haplotypes of a query diploid assembly and $R$ a reference haplotype. Three results are proved. **Theorem 1**: under a biallelic assumption, the counts of heterozygous and homozygous non-reference sites are determined uniquely by the pairwise Hamming distances among the three sequences; this is a combinatorial identity involving no population-genetic assumption. **Theorem 2**: if the three haplotypes are exchangeable, the ratio of the expected counts is exactly 2, proved from exchangeability alone. **Theorem 3**: in general the ratio is $2(1-F)/(1+F)$ with $F = 1 - \pi_w/\pi_b$; we prove that $F$ equals $F_{\mathrm{ST}}$ under a population split model and $F_{\mathrm{IS}}$ under the inbreeding model specified in Proposition 3, with $F \in (-1, 1]$ and $\bar{r}$ ranging over $[0, \infty)$. Section 7 provides two technical checks, and Section 8 summarizes the minimal assumptions on which each result depends. Throughout, the results concern biallelic single-nucleotide sites on a fixed common site set, and the ratio of expected counts rather than the expectation of the ratio.


#### 1. Notation and assumptions

**Definition 1 (data).** Let $\mathcal{S}$ be a finite set of sites with $|\mathcal{S}| = L$, fixed throughout. Three sequences are given on $\mathcal{S}$: the two haplotypes $Q_1, Q_2$ of a query diploid assembly, and a reference haplotype $R$. Each site of $\mathcal{S}$ is an aligned, callable single-nucleotide position at which all three sequences are assigned a state; indels, structural variants and sites of length other than one are excluded from $\mathcal{S}$ by construction. Positions at which all three sequences agree may remain in $\mathcal{S}$, because they contribute zero to every count and distance. For a sequence $X$ and a site $s \in \mathcal{S}$, write $X(s)$ for its base state. For two sequences $X, Y$ write

$$d(X, Y) = \#\{s \in \mathcal{S} : X(s) \neq Y(s)\}$$

for their Hamming distance.

**Definition 2 (genotype classes).** Classify each site by the state of $(Q_1, Q_2)$ relative to $R$:

$$\begin{aligned} \mathrm{HET} &= \{s : Q_1(s) \neq Q_2(s)\}, \\ \mathrm{HOM} &= \{s : Q_1(s) = Q_2(s) \neq R(s)\}, \\ \mathrm{REF} &= \{s : Q_1(s) = Q_2(s) = R(s)\}, \end{aligned}$$

and set $\#\mathbf{het} = |\mathrm{HET}|$, $\#\mathbf{hom} = |\mathrm{HOM}|$. The three sets are pairwise disjoint with union $\mathcal{S}$. When $\#\mathbf{hom} > 0$, the observed finite-site ratio is

$$r_{\mathrm{obs}} = \frac{\#\mathbf{het}}{\#\mathbf{hom}}.$$

The theoretical target introduced in Section 3 is the ratio of expected counts, $\bar{r} = \mathbb{E}[\#\mathbf{het}]/\mathbb{E}[\#\mathbf{hom}]$. These two quantities are distinct: $\bar{r}$ is not generally equal to $\mathbb{E}[r_{\mathrm{obs}}]$.

This classification corresponds to the phased dipcall output: $\mathrm{HET}$ to genotypes `1|0` or `0|1` (under the biallelic assumption of Definition 3), $\mathrm{HOM}$ to `1|1`, and $\mathrm{REF}$ to `0|0`. The proofs below use only Definition 2 and make no further reference to the VCF representation.

**Definition 3 (assumptions).**

- **(A1) Biallelic.** For each $s \in \mathcal{S}$, the states of the three sequences at $s$ lie in a two-element set $\{a_s, b_s\}$.
- **(A2) Common callable domain.** All three sequences are defined at every site of $\mathcal{S}$, and $\mathcal{S}$ is the same set for all three: no site is scored for one sequence and missing for another. In practice $\mathcal{S}$ is the callable region on which the comparison is performed.
- **(A3) No recurrent mutation (infinite sites).** At most one mutation per site on the genealogy of $Q_1, Q_2, R$.
- **(A4) Within-individual exchangeability.** The joint law of $(Q_1, Q_2, R)$ is invariant under the swap $Q_1 \leftrightarrow Q_2$.
- **(A5) Full exchangeability.** The joint law of $(Q_1, Q_2, R)$ is invariant under every permutation of the three labels.

Section 2 uses only (A1)–(A2); Section 3 adds (A4); Section 4 uses (A5). Assumption (A3) is used only where the counts are interpreted as numbers of mutations on a genealogy (Remark 3, Proposition 5); the combinatorial identity itself does not require it.

By (A1), at each site the three sequences take at most two distinct states, so the results below are statements about biallelic single-nucleotide sites only. Multiallelic sites, at which the three sequences take three distinct states, are treated separately in Remark 4; indels and structural variants lie outside $\mathcal{S}$ by Definition 1 and are not covered by any result in this note.

> **Convention 1 (the site set is part of the data)**
>
> *All quantities defined below — $\#\mathrm{het}$, $\#\mathrm{hom}$, $n_1, n_2, n_R$, $d(\cdot,\cdot)$, $\pi_w$, $\pi_b$, $F$ — are functions of the pair (sequences, $\mathcal{S}$). When several references $R^{(1)}, \ldots, R^{(k)}$ are compared, their comparison-specific callable sets must first be represented as homologous sites in one common coordinate system. All ratios must then be calculated on the same site set, for example $\mathcal{S} = \bigcap_j \mathcal{S}^{(j)}$, where $\mathcal{S}^{(j)}$ is the projected callable site set of comparison $j$.*

The reason is Corollary 1 below: $\#\mathrm{het}$ is a function of $Q_1, Q_2$ and $\mathcal{S}$ alone. If each comparison uses its own callable region $\mathcal{S}^{(j)}$, then $\#\mathrm{het}$ varies across comparisons through $\mathcal{S}^{(j)}$ even though it carries no information about $R^{(j)}$, and the resulting variation in $r_{\mathrm{obs}}$ mixes this artefact with the genuine dependence of $\#\mathrm{hom}$ on $R^{(j)}$. On a common $\mathcal{S}$, $\#\mathrm{het}$ is constant across $j$ and all variation in $r_{\mathrm{obs}}$ is attributable to $\#\mathrm{hom}$. Convention 1 is a condition on the analysis, not a probabilistic assumption, and is used implicitly wherever ratios from different references are compared.

#### 2. Combinatorial layer: a deterministic identity

**Definition 4 (separating counts).** For $x \in \{Q_1, Q_2, R\}$, let $n_x$ be the number of sites at which $x$ differs from both of the other two sequences:

$$n_1 = \#\{s : Q_1(s) \neq Q_2(s) = R(s)\},$$
$$n_2 = \#\{s : Q_2(s) \neq Q_1(s) = R(s)\},$$
$$n_R = \#\{s : R(s) \neq Q_1(s) = Q_2(s)\}.$$

Note that $n_1, n_2, n_R$ are deterministic functions of the three sequences; their definition invokes no ancestral state, no genealogy, and no model.

**Lemma 1 (completeness of the classification)**

*Under (A1)–(A2), $\mathcal{S}$ is partitioned into four pairwise disjoint subsets, with counts $|\mathrm{REF}|$, $n_1$, $n_2$, $n_R$. Moreover*

$$\#\mathrm{het} = n_1 + n_2, \qquad \#\mathrm{hom} = n_R.$$

**Proof.** Fix $s \in \mathcal{S}$. By (A1) the states of the three sequences at $s$ form a triple over the two-element set $\{a_s, b_s\}$. Three elements taking two values admit only two possibilities: all three agree, or exactly one differs from the other two. The former is $s \in \mathrm{REF}$; the latter splits into three mutually exclusive cases according to which sequence is the odd one out, with counts $n_1, n_2, n_R$. The four cases are exhaustive and mutually exclusive, proving the first claim.

Next, $s \in \mathrm{HET}$ iff $Q_1(s) \neq Q_2(s)$. By the classification above this happens exactly in the cases "$Q_1$ odd" and "$Q_2$ odd" (if $R$ is the odd one then $Q_1(s) = Q_2(s)$, and likewise if all three agree), so $\#\mathrm{het} = n_1 + n_2$. And $s \in \mathrm{HOM}$ iff $Q_1(s) = Q_2(s) \neq R(s)$, which is exactly the case "$R$ odd", so $\#\mathrm{hom} = n_R$. □

**Remark 1.** The second claim of Lemma 1 is the central observation of this note: $\mathrm{HOM}$ corresponds to the case "the reference is the odd one out", and *not* to "the query carries the derived allele". Since dipcall has no access to the ancestral state, the genotype classification depends only on the *unoriented* partition pattern of the three sequences, independently of which side is derived. Treating $R$ as an ancestral sequence would restrict $\mathrm{HOM}$ to the case "query homozygous derived" and thereby omit the sites at which the query is homozygous ancestral while the reference carries the derived allele; in Lemma 1 both are correctly merged into the single count $n_R$.

**Theorem 1 (three-haplotype identity)**

*Under (A1)–(A2),*

$$\#\mathrm{het} = d(Q_1, Q_2), \qquad \#\mathrm{hom} = \tfrac{1}{2}\left[\, d(Q_1, R) + d(Q_2, R) - d(Q_1, Q_2) \,\right],$$

*and hence, when $\#\mathrm{hom} > 0$,*

$$r_{\mathrm{obs}} = \frac{2\, d(Q_1, Q_2)}{d(Q_1, R) + d(Q_2, R) - d(Q_1, Q_2)}.$$

**Proof.** By the classification of Lemma 1, every non-$\mathrm{REF}$ site belongs to exactly one of $n_1, n_2, n_R$. Tabulating the contribution of each class to each of the three distances:

| site type | $d(Q_1, Q_2)$ | $d(Q_1, R)$ | $d(Q_2, R)$ |
|---|---|---|---|
| $Q_1$ odd (counted in $n_1$) | 1 | 1 | 0 |
| $Q_2$ odd (counted in $n_2$) | 1 | 0 | 1 |
| $R$ odd (counted in $n_R$) | 0 | 1 | 1 |
| all three agree | 0 | 0 | 0 |

Summing over sites gives

$$d(Q_1, Q_2) = n_1 + n_2, \qquad d(Q_1, R) = n_1 + n_R, \qquad d(Q_2, R) = n_2 + n_R. \tag{2.1}$$

The first, combined with Lemma 1, yields $\#\mathrm{het} = n_1 + n_2 = d(Q_1, Q_2)$. Adding the last two equations of (2.1) and subtracting the first,

$$d(Q_1, R) + d(Q_2, R) - d(Q_1, Q_2) = (n_1 + n_R) + (n_2 + n_R) - (n_1 + n_2) = 2n_R,$$

and $\#\mathrm{hom} = n_R$ from Lemma 1 gives the second equation. The expression for $r_{\mathrm{obs}}$ is their quotient. □

**Remark 2 (linear invertibility).** (2.1) is a linear map from $(n_1, n_2, n_R)$ to the three distances, with matrix

$$M = \begin{pmatrix} 1 & 1 & 0 \\ 1 & 0 & 1 \\ 0 & 1 & 1 \end{pmatrix}, \qquad \det M = -2 \neq 0.$$

Since $M$ is non-singular the map is a bijection, and Theorem 1 is the third row of $M^{-1}$. Precisely: for a fixed site set $\mathcal{S}$ satisfying (A1)–(A2), the vector $(n_1, n_2, n_R)$ and the vector $\big(d(Q_1, Q_2), d(Q_1, R), d(Q_2, R)\big)$ determine each other uniquely. The equivalence is between these two triples of integers and nothing more: neither determines $|\mathrm{REF}|$, $L$, the identity of the sites involved, or the sequences themselves. In particular $\#\mathrm{het}$ and $\#\mathrm{hom}$ can be computed from the three distances, and conversely, but neither triple identifies which sites are variant.

**Remark 3 (tree-metric reading).** Under (A3), $n_1, n_2, n_R$ are the numbers of mutations on the three pendant edges of the unrooted three-leaf tree, and the second equation of Theorem 1 is the Buneman three-point formula: $n_R$ is the length of the path from $R$ to the internal node joining $Q_1$ and $Q_2$. Note however that the proof of Theorem 1 does *not* use (A3): it is purely combinatorial and holds for any three sequences satisfying (A1)–(A2), whether or not they were generated by a tree-like history.

**Corollary 1 (the het count does not involve the reference)**

*Under (A1)–(A2),* $\#\mathrm{het} = d(Q_1, Q_2)$ *is a function of* $Q_1, Q_2$ *alone and does not depend on* $R$*.*

**Proof.** By the first equation of Theorem 1, the right-hand side does not involve $R$. □

Corollary 1 concerns the idealised data of Definition 1, in which all three sequences are observed without error. In practice, alignment and variant-calling error rates can themselves vary with the reference, so the measured $\#\mathrm{het}$ can retain a reference-dependent component introduced by the measurement process. Convention 1 removes dependence caused by differences in the callable site set, but it does not remove this source of measurement error.

**Remark 4 (correction when (A1) fails).** Suppose at some sites the three sequences take three distinct states, violating (A1) (dipcall records these as `1|2` or `2|1`), and let $m$ be the number of such sites. Each contributes 1 to all three distances, so (2.1) becomes $d(Q_1, Q_2) = n_1 + n_2 + m$ and likewise for the others, whence

$$\#\mathrm{het} = n_1 + n_2 + m = d(Q_1, Q_2), \qquad \#\mathrm{hom} = n_R = \tfrac{1}{2}\big[d(Q_1, R) + d(Q_2, R) - d(Q_1, Q_2)\big] - \tfrac{m}{2}.$$

The first equation of Theorem 1 thus remains exact (such sites satisfy $Q_1 \neq Q_2$ and so lie in $\mathrm{HET}$), while the second acquires a correction $-m/2$. Assumption (A1) is therefore load-bearing only in the second equation.

Consequently, the genotype-based ratio and the biallelic distance expression diverge when $m > 0$. Provided $n_R > 0$, the genotype-based ratio is $(n_1 + n_2 + m)/n_R$, whereas the distance expression in Theorem 1 gives

$$\frac{2\,d(Q_1, Q_2)}{d(Q_1, R) + d(Q_2, R) - d(Q_1, Q_2)} = \frac{2(n_1 + n_2 + m)}{2n_R + m} = \frac{n_1 + n_2 + m}{n_R}\left(1 + \frac{m}{2n_R}\right)^{-1}.$$

The distance expression therefore underestimates the genotype-based ratio by the factor $(1+m/(2n_R))^{-1}$. Multiallelic sites also increase the genotype-based ratio relative to the biallelic-only value $(n_1+n_2)/n_R$, because they enter its numerator but not its denominator. For $m/n_R \ll 1$, both effects are of order $m/n_R$ and vanish on a strictly biallelic site set.

#### 3. Expectation layer

From this section on, $(Q_1, Q_2, R)$ is treated as random, i.e. drawn from some sampling model.

**Definition 5.** Under (A4) set

$$\pi_w = \mathbb{E}\big[d(Q_1, Q_2)\big], \qquad \pi_b = \mathbb{E}\big[d(Q_1, R)\big] = \mathbb{E}\big[d(Q_2, R)\big].$$

The second equality is guaranteed by (A4) (the law is unchanged by swapping $Q_1 \leftrightarrow Q_2$). Thus, $\pi_w$ is the expected number of heterozygous sites in the query and $\pi_b$ is the expected number of query–reference differences, both counted over $\mathcal{S}$. Dividing both quantities by $L$ gives their per-site versions without changing $F$ or $\bar{r}$.

**Proposition 1 (expected counts)**

*Under (A1), (A2), (A4),*

$$\mathbb{E}[\#\mathrm{het}] = \pi_w, \qquad \mathbb{E}[\#\mathrm{hom}] = \pi_b - \tfrac{1}{2}\pi_w.$$

**Proof.** Take expectations in the two equations of Theorem 1 and use linearity together with Definition 5:

$$\mathbb{E}[\#\mathrm{het}] = \mathbb{E}[d(Q_1, Q_2)] = \pi_w,$$

$$\mathbb{E}[\#\mathrm{hom}] = \tfrac{1}{2}\big(\mathbb{E}[d(Q_1, R)] + \mathbb{E}[d(Q_2, R)] - \mathbb{E}[d(Q_1, Q_2)]\big) = \tfrac{1}{2}(2\pi_b - \pi_w) = \pi_b - \tfrac{1}{2}\pi_w. \qquad \square$$

Write the ratio of expectations as

$$\bar{r} = \frac{\mathbb{E}[\#\mathrm{het}]}{\mathbb{E}[\#\mathrm{hom}]}$$

(All results below concern $\bar{r}$, which is not generally equal to $\mathbb{E}[r_{\mathrm{obs}}]$. This note does not establish a finite-site concentration result for $r_{\mathrm{obs}}$; treating it as an estimator of $\bar{r}$ requires a separate uncertainty or concentration analysis.)

#### 4. The exchangeable case

**Theorem 2**

*Under (A1), (A2), (A5), if* $\mathbb{E}[\#\mathrm{hom}] > 0$ *then*

$$\bar{r} = 2 \qquad \text{exactly.}$$

**Proof.** By Definition 4, $n_1, n_2, n_R$ are one and the same function applied at different labels: writing $N(x; X, Y, Z)$ for the number of sites at which $x$ differs from both other members of the triple $(X, Y, Z)$, we have

$$n_1 = N(Q_1; Q_1, Q_2, R), \quad n_2 = N(Q_2; Q_1, Q_2, R), \quad n_R = N(R; Q_1, Q_2, R),$$

and $N$ is symmetric in the order of its latter two arguments. Applying the permutation that exchanges the first and third labels, (A5) gives that $(R, Q_2, Q_1)$ has the same law as $(Q_1, Q_2, R)$, hence

$$n_R = N(R; R, Q_2, Q_1) \overset{d}{=} N(Q_1; Q_1, Q_2, R) = n_1.$$

Similarly $n_R \overset{d}{=} n_2$. The three counts are therefore identically distributed and in particular have equal means:

$$\mathbb{E}[n_1] = \mathbb{E}[n_2] = \mathbb{E}[n_R] =: \nu.$$

By Lemma 1, $\mathbb{E}[\#\text{het}] = 2\nu$ and $\mathbb{E}[\#\text{hom}] = \nu$, so $\bar{r} = 2$. □

**Interpretation.** Under the biallelic classification, HET pools the two mutually exclusive cases in which $Q_1$ or $Q_2$ is the unique sequence state, whereas HOM contains the single case in which $R$ is unique. Full exchangeability makes the expected counts of these three odd-one-out classes equal; HET therefore contains two equal expected contributions and HOM one, giving the $2:1$ ratio.

**Remark 5.** The proof uses only exchangeability and the combinatorial classification of Lemma 1; no mutation model, coalescent, frequency spectrum, or demographic history enters. The conclusion therefore holds automatically for any neutral sampling model that makes $(Q_1, Q_2, R)$ exchangeable, including arbitrary time-varying population size $N(t)$, arbitrary Cannings models, and $\Lambda$-coalescents.

Two directions should be distinguished. Since (A5) is a hypothesis of Theorem 2, the conclusion $\bar{r} = 2$ can be obtained by this route only if (A5) holds; (A5) fails whenever the sampling scheme does not treat the three labels alike, for instance when $R$ is drawn from a different population than $Q_1, Q_2$ (Proposition 2) or when the two query haplotypes are correlated by inbreeding (Proposition 3), and these are examples rather than an exhaustive list. But failure of (A5) does not imply $\bar{r} \neq 2$: by Corollary 2 below the exact condition is $\pi_w = \pi_b$, which is a statement about two first moments and can hold under non-exchangeable models. (A5) is sufficient, not necessary.

Two model-specific checks of Theorem 2 are collected in Section 7 so that the general result and its population-genetic interpretations remain on the main line of the argument.

#### 5. The general case

**Definition 6.** Assume $\pi_b > 0$ and set

$$F = 1 - \frac{\pi_w}{\pi_b}.$$

The assumption $\pi_b > 0$ is needed for $F$ to be defined; it fails only if $Q_i$ and $R$ are almost surely identical on $\mathcal{S}$, in which case $\pi_w = 0$ as well by the triangle inequality and both counts vanish.

**Theorem 3**

*Under (A1), (A2), (A4), if* $\mathbb{E}[\#\text{hom}] > 0$ *then*

$$\bar{r} = \frac{2\pi_w}{2\pi_b - \pi_w} = \frac{2(1-F)}{1+F}.$$

*Moreover* $F \in (-1, 1]$*, and on that interval* $F \mapsto \bar{r}$ *is a strictly decreasing bijection onto* $[0, \infty)$*, with* $\bar{r} = 2 \iff F = 0$*.*

**Proof.** By Proposition 1,

$$\bar{r} = \frac{\pi_w}{\pi_b - \pi_w/2} = \frac{2\pi_w}{2\pi_b - \pi_w}.$$

Substituting $\pi_w = (1-F)\pi_b$,

$$\bar{r} = \frac{2(1-F)\pi_b}{2\pi_b - (1-F)\pi_b} = \frac{2(1-F)}{1+F},$$

the positivity of the denominator $1+F$ being established next.

**Range.** *Upper endpoint, attained.* Since $\pi_w = \mathbb{E}[d(Q_1, Q_2)] \geq 0$ and $\pi_b > 0$ we get $F \leq 1$, with $F = 1$ iff $\pi_w = 0$. The value $F = 1$ is admissible: it means the query is almost surely homozygous throughout $\mathcal{S}$, so $\mathbb{E}[\#\text{het}] = 0$ while $\mathbb{E}[\#\text{hom}] = \pi_b > 0$, the hypothesis $\mathbb{E}[\#\text{hom}] > 0$ holds, and $\bar{r} = 0$. Nothing is degenerate; $\bar{r} = 0$ is a legitimate value of the ratio of expectations. Hence $F = 1$ is included.

*Lower endpoint, excluded.* The Hamming distance satisfies the triangle inequality $d(Q_1, Q_2) \leq d(Q_1, R) + d(Q_2, R)$ site by site, so it survives taking expectations and gives $\pi_w \leq 2\pi_b$, i.e. $F \geq -1$. The endpoint is removed by the hypothesis: by Proposition 1,

$$\mathbb{E}[\#\text{hom}] = \pi_b - \tfrac{1}{2}\pi_w = \tfrac{1}{2}(2\pi_b - \pi_w) = \tfrac{1}{2}\pi_b(1+F),$$

so, $\pi_b$ being positive,

$$\mathbb{E}[\#\text{hom}] > 0 \iff 1 + F > 0 \iff F > -1.$$

Thus $F = -1$ is exactly the case $\mathbb{E}[\#\text{hom}] = 0$ excluded by hypothesis, and $F \in (-1, 1]$. Note that the same computation shows $\mathbb{E}[\#\text{hom}] \geq 0$ unconditionally, as befits a count, and that the denominator $1+F$ in the statement is strictly positive, which justifies the division performed above.

**Monotonicity and range of $\bar{r}$.** On $F \in (-1, 1]$,

$$\frac{d}{dF}\left[\frac{2(1-F)}{1+F}\right] = \frac{-2(1+F) - 2(1-F)}{(1+F)^2} = \frac{-4}{(1+F)^2} < 0,$$

so $\bar{r}$ is strictly decreasing, hence injective. It is continuous on $(-1, 1]$ with $\bar{r} = 0$ at $F = 1$ and $\bar{r} \to +\infty$ as $F \downarrow -1$, so by the intermediate value theorem its image is exactly $[0, \infty)$; the map is a bijection $(-1, 1] \to [0, \infty)$. Finally $\bar{r} = 2 \iff 2(1-F) = 2(1+F) \iff F = 0$. □

**Reference-distance interpretation.** When the within-query expectation $\pi_w$ is fixed, increasing the query-to-reference expectation $\pi_b$ increases $F = 1 - \pi_w/\pi_b$ and therefore decreases $\bar{r}$. This monotonic interpretation is conditional on using the same homologous callable site set across reference comparisons; Proposition 2 gives its explicit population-split form.

**Corollary 2 (necessary and sufficient condition for the ratio 2)**

*Under (A1), (A2), (A4),* $\bar{r} = 2 \iff \pi_w = \pi_b$.

**Proof.** By Theorem 3, $\bar{r} = 2 \iff F = 0 \iff \pi_w = \pi_b$. □

**Remark 6.** Relation to Theorem 2: (A5) implies $\pi_w = \pi_b$, since under (A5) $d(Q_1, Q_2) \overset{d}{=} d(Q_1, R)$. Theorem 2 is therefore a special case of Corollary 2. But $\pi_w = \pi_b$ is strictly weaker than (A5): it asks only that two first moments agree, not that the joint law be exchangeable. The necessary and sufficient condition for $\bar{r} = 2$ is thus this first-moment condition, not exchangeability.

**Remark 7 (inversion).** By Theorem 3 the map $F \mapsto \bar{r}$ is a strictly decreasing bijection $(-1, 1] \to [0, \infty)$, so it is invertible on the whole of $[0, \infty)$:

$$F = \frac{2 - \bar{r}}{2 + \bar{r}} \qquad (\bar{r} \geq 0),$$

the denominator being positive throughout. Thus $\bar{r}$ determines $F$ uniquely, and $\bar{r} = 0$ returns $F = 1$ as it should.

The ratio of the two summary statistics can also be recovered, but only when it is defined. Dividing Proposition 1 through,

$$\frac{\pi_b}{\pi_w} = \frac{1}{2} + \frac{1}{\bar{r}} \qquad (\bar{r} > 0),$$

which requires $\bar{r} > 0$, equivalently $\pi_w > 0$, equivalently $F < 1$. At $\bar{r} = 0$ we have $\pi_w = 0$ and the quotient $\pi_b/\pi_w$ is not defined, while $F = 1$ still is; the form $F = (2 - \bar{r})/(2 + \bar{r})$ is therefore the one to use in general. Finally $\bar{r} > 2 \iff F < 0 \iff \pi_w > \pi_b$, and $\bar{r} < 2 \iff F > 0 \iff \pi_w < \pi_b$.

#### 6. Two model readings of $F$

The $F$ of Definition 6 is given purely by $\pi_w$ and $\pi_b$ and carries no model content of its own. The following two propositions identify it with the classical $F_{\mathrm{ST}}$ and $F_{\mathrm{IS}}$ within a population split model and an inbreeding model respectively. The coalescent time scale and mutation parameterisation used in the split-model calculation are detailed in Section 7.

**Proposition 2 (split model: $F = F_{\mathrm{ST}}$)**

*Let two populations, each of size $N$, diverge at time $T$ (in units of $2N$ generations) from an ancestral population also of size $N$, with no subsequent migration; let $Q_1, Q_2$ be drawn from population $P$ and $R$ from $P'$. Then under the neutral infinite-sites model*

$$\pi_w = \theta, \qquad \pi_b = \theta(T+1), \qquad F = \frac{T}{T+1} = F_{\mathrm{ST}}, \qquad \bar{r} = \frac{2}{1+2T},$$

*where $F_{\mathrm{ST}}$ is taken in the Hudson–Slatkin–Maddison sense, $F_{\mathrm{ST}} = 1 - \pi_w/\pi_b$ with $\pi_w, \pi_b$ the expected numbers of differences for a within- and a between-population pair.*

**Proof.** Under the infinite-sites model, two lineages coalescing at time $t$ differ at $\theta t$ sites in expectation (total branch length $2t$, mutation rate $\theta/2$ per unit length). It therefore suffices to obtain the expected coalescence time for each type of pair.

*Within-population pair.* $Q_1, Q_2$ are both in $P$. Before $T$ they coalesce at rate 1 within $P$, which has size $N$; if they have not, at time $T$ they enter the ancestral population, also of size $N$, and continue to coalesce at rate 1. The rate is 1 throughout, so $t_{12} \sim \mathrm{Exp}(1)$, $\mathbb{E}[t_{12}] = 1$, and

$$\pi_w = \theta\,\mathbb{E}[t_{12}] = \theta.$$

This is independent of $T$, consistent with Corollary 1: $\#\mathrm{het}$ does not depend on $R$, so a fortiori not on the divergence time between $P$ and the population from which $R$ is drawn.

*Between-population pair.* $Q_i$ and $R$ lie in two populations exchanging no migrants and so cannot coalesce before $T$; at time $T$ both lineages enter the ancestral population and thereafter coalesce at rate 1. Hence $t_{iR} = T + E$ with $E \sim \mathrm{Exp}(1)$, $\mathbb{E}[t_{iR}] = T + 1$, and

$$\pi_b = \theta\,\mathbb{E}[t_{iR}] = \theta(T+1).$$

Definition 6 gives $F = 1 - \theta/[\theta(T+1)] = T/(T+1)$, which is exactly $F_{\mathrm{ST}} = 1 - \pi_w/\pi_b$. Theorem 3 then gives

$$\bar{r} = \frac{2(1 - \frac{T}{T+1})}{1 + \frac{T}{T+1}} = \frac{2 \cdot \frac{1}{T+1}}{\frac{2T+1}{T+1}} = \frac{2}{2T+1}. \qquad \square$$

**Remark 8 (branch-length check).** One can verify this without $F$. Without migration, $R$ cannot coalesce with either $Q_i$ first, so the topology is necessarily $((Q_1, Q_2), R)$. By the tree-metric form of (2.1), $d(Q_i, R) = L_{Q_i} + L_R$ and $d(Q_1, Q_2) = L_{Q_1} + L_{Q_2}$, so

$$\mathbb{E}[L_{Q_1} + L_{Q_2}] = 2\,\mathbb{E}[t_{12}] = 2, \qquad \mathbb{E}[L_R] = 2\,\mathbb{E}[t_{iR}] - \mathbb{E}[t_{12}] = 2(T+1) - 1 = 2T+1,$$

giving $\bar{r} = 2/(2T+1)$ in agreement with Proposition 2.

**Proposition 3 (inbreeding model: $F = F_{\mathrm{IS}}$)**

*Let the query individual have inbreeding coefficient* $F_{\mathrm{IS}} \in [0,1]$ *relative to its population, and let* $R$ *be a haplotype drawn at random from that same population, independently of the query individual. Then at every site with frequency* $p \in (0,1)$*,*

$$\frac{\Pr(\mathrm{HET})}{\Pr(\mathrm{HOM})} = \frac{2(1 - F_{\mathrm{IS}})}{1 + F_{\mathrm{IS}}},$$

*hence, provided* $\sum_s p_s(1 - p_s) > 0$*,* $\bar{r} = 2(1 - F_{\mathrm{IS}})/(1 + F_{\mathrm{IS}})$ *and* $F = F_{\mathrm{IS}}$*.*

The restriction $F_{\mathrm{IS}} \in [0,1]$ guarantees that the genotype probabilities used below are non-negative for every allele frequency. If negative $F_{\mathrm{IS}}$ values are to be included, they must additionally satisfy, at every analysed site, $F_{\mathrm{IS}} \geq -\min\{p/(1-p), (1-p)/p\}$; that extension is not required for the inbreeding model considered here.

**Proof.** Fix a site, let $p$ be the population frequency of the derived allele, and write $F = F_{\mathrm{IS}}$. By definition of the inbreeding coefficient the individual's genotype frequencies are

$$\Pr(Q_1 \neq Q_2) = 2p(1-p)(1-F),$$

$$\Pr(Q_1 = Q_2 = \text{derived}) = p^2 + Fp(1-p), \qquad \Pr(Q_1 = Q_2 = \text{ancestral}) = (1-p)^2 + Fp(1-p),$$

which sum to $p^2 + 2p(1-p) + (1-p)^2 = 1$, as required. $R$ is drawn independently and is derived with probability $p$. Hence

$$\begin{aligned}
\Pr(\mathrm{HOM}) &= \Pr(Q_1 = Q_2 = \text{derived})\,(1-p) + \Pr(Q_1 = Q_2 = \text{ancestral})\,p \\
&= \big[p^2 + Fp(1-p)\big](1-p) + \big[(1-p)^2 + Fp(1-p)\big]p \\
&= \underbrace{p^2(1-p) + (1-p)^2 p}_{=\,p(1-p)} + Fp(1-p)\big[(1-p) + p\big] \\
&= p(1-p)(1+F).
\end{aligned}$$

For $p \in (0,1)$ we have $p(1-p) > 0$, and because $F \in [0,1]$, $\Pr(\mathrm{HOM}) = p(1-p)(1+F) > 0$; dividing into $\Pr(\mathrm{HET}) = 2p(1-p)(1-F)$ gives the stated ratio, which is independent of $p$. At a site with $p \in \{0,1\}$ both probabilities are 0 and the site-wise ratio is undefined, but such a site contributes nothing to either count.

Finally, to check $F = F_{\text{IS}}$, summing site-wise gives

$$\pi_w = \sum_s 2p_s(1-p_s)(1-F), \qquad \pi_b = \sum_s \Pr\big(Q_i(s) \neq R(s)\big) = \sum_s 2p_s(1-p_s),$$

the expression for $\pi_b$ holding because a single allele of $Q_i$ and the independently drawn $R$ are each marginally derived with probability $p_s$ and are independent of one another (inbreeding affects the correlation between the individual's two alleles, not the marginal law of one allele). Writing $V = \sum_s 2p_s(1-p_s)$, the hypothesis $\sum_s p_s(1-p_s) > 0$ says $V > 0$, so $\pi_b = V > 0$ as Definition 6 requires, and

$$1 - \frac{\pi_w}{\pi_b} = 1 - \frac{V(1-F)}{V} = F = F_{\text{IS}}.$$

Theorem 3 then returns $\bar{r} = 2(1-F_{\text{IS}})/(1+F_{\text{IS}})$, consistent with the site-wise computation. □

**Remark 9.** Propositions 2 and 3 yield the same functional form $2(1-F)/(1+F)$, with $F$ defined in both models by Definition 6. The same value of $\bar{r}$ can therefore arise from different mechanisms. Without additional information, $\bar{r}$ alone cannot determine whether a departure from 2 reflects between-population divergence, inbreeding or another process affecting $\pi_w/\pi_b$.

#### 7. Technical checks and extensions

The following calculations strengthen or verify Theorem 2 within specific models. Neither calculation is required for the deterministic identity, the expectation formula or the general transformation in Theorem 3.

**Proposition 4 (site-wise strengthening)**

*Suppose that at each site $s$ there is a frequency $p_s \in [0,1]$ such that $Q_1(s), Q_2(s), R(s)$ are i.i.d., each derived with probability $p_s$. Then for every $s$ with $p_s \in (0,1)$,*

$$\frac{\Pr(s \in \text{HET})}{\Pr(s \in \text{HOM})} = 2,$$

*and consequently $\bar{r} = 2$ for any $\{p_s\}$, deterministic or random, provided $\sum_s \mathbb{E}[p_s(1-p_s)] > 0$.*

**Proof.** Fix $s$ and drop the subscript, writing $p$. From three independent draws,

$$\Pr(\text{HET}) = \Pr(Q_1 \neq Q_2) = 2p(1-p).$$

HOM requires $Q_1 = Q_2 \neq R$, which splits into two mutually exclusive cases according to the shared value of $Q_1 = Q_2$:

$$\Pr(\text{HOM}) = \underbrace{p^2(1-p)}_{Q_1=Q_2=\text{derived},\, R=\text{ancestral}} + \underbrace{(1-p)^2 p}_{Q_1=Q_2=\text{ancestral},\, R=\text{derived}} = p(1-p)\big[p + (1-p)\big] = p(1-p).$$

If $p \in (0,1)$, then $p(1-p) > 0$ and the ratio is $2p(1-p)/[p(1-p)] = 2$, independent of $p$. If $p \in \{0,1\}$, both probabilities vanish and the site-wise ratio is undefined; such a site contributes 0 to both counts. Summing over $s$ , and taking expectations first if the $p_s$ are random, gives

$$\mathbb{E}[\#\text{het}] = \sum_s 2\,\mathbb{E}[p_s(1-p_s)], \qquad \mathbb{E}[\#\text{hom}] = \sum_s \mathbb{E}[p_s(1-p_s)].$$

If $\sum_s \mathbb{E}[p_s(1-p_s)] > 0$, the denominator is non-zero and the quotient is 2. If the sum vanishes, every site is monomorphic almost surely, both expected counts are 0 and $\bar{r}$ is undefined. □

**Remark 10.** Proposition 4 shows that the result does not depend on the distribution of allele frequencies. The ratio is already 2 at each polymorphic $p$, so any weighting across sites preserves it. This is the quantitative counterpart of Remark 1: HOM represents the reference being the odd sequence state and is independent of polarisation.

**Background (the Kingman coalescent and the infinite-sites model).** Proposition 5 and the split-model calculation in Proposition 2 use the following standard neutral-genealogy conventions. Consider a population of $N$ diploid individuals ($2N$ haplotypes) reproducing neutrally, and sample $n$ haplotypes at the present. Trace their ancestral lineages backwards in time, measuring time in units of $2N$ generations. In the limit $N \to \infty$, Kingman's coalescent states that while $k$ lineages remain, each of the $\binom{k}{2}$ pairs coalesces at rate 1, so the waiting time until the next coalescence is

$$T_k \sim \mathrm{Exp}\left(\binom{k}{2}\right), \qquad \mathbb{E}[T_k] = \binom{k}{2}^{-1}.$$

The $T_k$ are independent, and the pair that coalesces is chosen uniformly among the $\binom{k}{2}$ pairs, independently of all the $T_j$. Running from $k = n$ down to $k = 1$ produces the genealogy of the sample. For $n = 2$, the single waiting time is $T_2 \sim \mathrm{Exp}(1)$, so the coalescence time of a pair has mean 1. The construction is also exchangeable in the sample labels. For $n = 3$, each first-coalescing pair, and hence each unrooted topology, has probability $1/3$, independently of $T_3$ and $T_2$.

Under the infinite-sites model, mutations arise along the genealogy as a Poisson process of intensity $\theta/2$ per unit branch length, where $\theta = 4N\mu$ and $\mu$ is the per-sequence per-generation mutation rate. Each mutation occurs at a site not previously mutated. Thus, a branch of length $\ell$ carries $\theta\ell/2$ mutations in expectation, and two lineages coalescing at time $t$ differ at $\theta t$ sites in expectation. The expected total number of segregating sites is $\mathbb{E}[S_n] = \theta H_{n-1}$, where $H_m = \sum_{i=1}^{m} 1/i$.

**Proposition 5 (check under the Kingman coalescent)**

*Under the neutral Kingman coalescent with constant size $N$ and the infinite-sites mutation model ($\theta = 4N\mu$, time in units of $2N$ generations),*

$$\mathbb{E}[\#\mathrm{het}] = \theta, \qquad \mathbb{E}[\#\mathrm{hom}] = \theta/2, \qquad \bar{r} = 2.$$

**Proof.** Under (A3), interpret $n_x$ as the number of mutations on the pendant edge of $x$ in the unrooted three-leaf tree. A mutation on this edge is carried by $x$ alone and is therefore counted by $n_x$ in Definition 4. By the Poisson intensity $\theta/2$, $\mathbb{E}[n_x] = \frac{\theta}{2}\mathbb{E}[L_x]$, where $L_x$ is the length of that pendant edge.

Let $T_3, T_2$ be the durations with three and two lineages present, so $\mathbb{E}[T_3] = 1/3$ and $\mathbb{E}[T_2] = 1$. If $Q_1, Q_2$ coalesce first, with topology $((Q_1, Q_2), R)$, then

$$L_{Q_1} = L_{Q_2} = T_3, \qquad L_R = T_3 + 2T_2.$$

The latter is the sum of $R$'s external branch and the edge above the common ancestor of $Q_1, Q_2$. Mutations on either edge produce the same partition pattern and therefore contribute to $n_R$. The total $L_{Q_1} + L_{Q_2} + L_R = 3T_3 + 2T_2$ equals the total branch length of the coalescent.

Each topology occurs with probability $1/3$, independently of $T_3, T_2$, so

$$\mathbb{E}[L_R] = \tfrac{1}{3}\big(\mathbb{E}[T_3] + 2\mathbb{E}[T_2]\big) + \tfrac{2}{3}\,\mathbb{E}[T_3] = \mathbb{E}[T_3] + \tfrac{2}{3}\mathbb{E}[T_2] = 1.$$

By symmetry, $\mathbb{E}[L_{Q_1}] = \mathbb{E}[L_{Q_2}] = 1$. Hence $\mathbb{E}[n_x] = \theta/2$ for all three labels, and Lemma 1 gives

$$\mathbb{E}[\#\mathrm{het}] = 2 \cdot \tfrac{\theta}{2} = \theta, \qquad \mathbb{E}[\#\mathrm{hom}] = \tfrac{\theta}{2}.$$

As a consistency check, summing the three classes gives $\mathbb{E}[S_3] = 3\theta/2 = \theta H_2$, in agreement with Watterson's formula. □

**Remark 11.** Proposition 5 is consistent with Theorem 2 but strictly weaker. It additionally assumes (A3), constant population size and the Kingman model, none of which Theorem 2 requires.

#### 8. Summary of dependencies

| Result | Statement | Assumptions used |
|---|---|---|
| Lemma 1 | the four classes are exhaustive; $\#\mathrm{het} = n_1 + n_2$, $\#\mathrm{hom} = n_R$ | (A1), (A2) |
| Theorem 1 | counts = combinatorial identity in the pairwise distances | (A1), (A2) |
| Corollary 1 | $\#\mathrm{het}$ does not depend on $R$ | (A1), (A2) |
| Proposition 1 | $\mathbb{E}[\#\mathrm{het}] = \pi_w$, $\mathbb{E}[\#\mathrm{hom}] = \pi_b - \pi_w/2$ | (A1), (A2), (A4) |
| Theorem 3 | $\bar{r} = 2(1-F)/(1+F)$; $F \in (-1,1]$; strictly decreasing bijection onto $[0,\infty)$ | (A1), (A2), (A4) |
| Corollary 2 | $\bar{r} = 2 \iff \pi_w = \pi_b$ | (A1), (A2), (A4) |
| Theorem 2 | $\bar{r} = 2$ | (A1), (A2), (A5) |
| Proposition 2 | $F = F_{\mathrm{ST}} = T/(T+1)$ | (A1)–(A4) + split model |
| Proposition 3 | $F = F_{\mathrm{IS}}$ | (A1), (A2), (A4) + inbreeding model with $F_{\mathrm{IS}} \in [0,1]$ |
| Proposition 4 | the site-wise ratio equals 2 | (A1), (A2) + i.i.d. sampling within each site, $p_s \in (0,1)$ |
| Proposition 5 | $\mathbb{E}[\#\mathrm{het}] = \theta$, $\mathbb{E}[\#\mathrm{hom}] = \theta/2$ | (A1)–(A3) + constant-size Kingman |

The main chain is

$$\text{Lemma 1} \implies \text{Theorem 1} \implies \text{Proposition 1} \implies \text{Theorem 3} \implies \text{Corollary 2},$$

which uses only (A1), (A2), (A4). Theorem 2 is the case $F = 0$ and admits an independent proof from (A5) (Section 4). Assumption (A3) lies off the main chain; it serves only to read $n_x$ as a count of mutations on a genealogy (Remark 3, Remark 8 and Proposition 5).

The necessary and sufficient condition for $\bar{r} = 2$ is the first-moment condition $\pi_w = \pi_b$ (Corollary 2); exchangeability (A5) is one sufficient condition for it.

**Scope.** Every statement above concerns biallelic single-nucleotide sites on a fixed common site set (Definition 1, (A1)–(A2), Convention 1). Multiallelic sites are covered only by the correction term of Remark 4; indels and structural variants are outside $\mathcal{S}$ and are not covered. All results concern the ratio of expectations $\bar{r}$, not $\mathbb{E}[r_{\mathrm{obs}}]$ or the realised finite-site ratio $r_{\mathrm{obs}}$ (Sections 1 and 3), and all hold for arbitrary $L$ with no appeal to asymptotics.
